# Skeletal muscle regeneration is altered in the R6/2 mouse model of Huntington’s disease

**DOI:** 10.1101/2022.01.11.475914

**Authors:** Sanzana Hoque, Marie Sjögren, Valérie Allamand, Kinga Gawlik, Naomi Franke, Madeleine Durbeej, Maria Björkqvist, Rana Soylu-Kucharz

**Affiliations:** Wallenberg Neuroscience Center, Brain Disease Biomarker Unit, Department of Experimental Medical Science, Lund University, Lund, Sweden; Muscle Biology Unit, Department of Experimental Medical Science, BMC C12, Lund University, 22184, Lund, Sweden; Sorbonne Université-Inserm, Centre de Recherche en Myologie, Institut de Myologie, F-75013 Paris, France

**Keywords:** Huntington’s disease, R6/2, satellite cells, muscle regeneration, inflammation

## Abstract

Huntington’s disease (HD) is caused by CAG repeat expansion in the huntingtin (*HTT*) gene. Skeletal muscle wasting alongside central pathology is a well-recognized phenomenon seen in patients with HD and HD mouse models. HD muscle atrophy progresses with disease and affects prognosis and quality of life. Satellite cells, progenitors of mature skeletal muscle fibers, are essential for proliferation, differentiation, and repair of muscle tissue in response to muscle injury or exercise.

In this study, we aim to investigate the effect of mutant HTT on the differentiation and regeneration capacity of HD muscle by employing *in vitro* mononuclear skeletal muscle cell isolation and *in vivo* acute muscle damage model in R6/2 mice.

We found that, similar to R6/2 adult mice, neonatal R6/2 mice also exhibit a significant reduction in myofiber width and morphological changes in gastrocnemius and soleus muscles compared to WT mice. Cardiotoxin (CTX)-induced acute muscle damage in R6/2 and WT mice showed that the Pax7+ satellite cell pool was dampened in R6/2 mice at 4 weeks post-injection, and R6/2 mice exhibited an altered inflammatory profile in response to acute damage.

Our results suggest that, in addition to the mutant HTT degenerative effects in mature muscle fibers, expression of mutant HTT in satellite cells might alter developmental and regenerative processes to contribute to the progressive muscle mass loss in HD. Taken together, the results presented here encourage further studies evaluating the underlying mechanisms of satellite cell dysfunction in HD mouse models.

## Introduction

Huntington’s disease (HD) is an inherited neurodegenerative disorder that is accompanied by skeletal muscle wasting alongside neurodegeneration. HD is caused by an expansion in the CAG triplet repeat in the gene encoding huntingtin (*HTT*) (Pagani et al., HDCRG, 1993) and is associated with progressive motor, cognitive and functional decline (Novak and Tabrizi, 2010, Walker, 2007). The mutated HTT protein (mHTT) causes the protein to gain toxic functions and aggregate, resulting in neuronal dysfunction and death (Bates et al., 2015, Ross and Tabrizi, 2011). In addition to central pathology (Novak and Tabrizi, 2010), altered energy metabolism, including weight loss and muscle atrophy, is seen in HD patients and HD mouse models (Carroll et al., 2015, Sathasivam et al., 1999, van der Burg et al., 2009, van der Burg et al., 2011, Bozzi and Sciandra, 2020). Both HD patients and HD transgenic mouse models exhibit motor deficits and muscle wasting even in the pre-symptomatic stage of HD (Busse et al., 2008, Kosinski et al., 2007, Trejo et al., 2004, Zielonka et al., 2014). The underlying mechanisms of muscle wasting in HD are still not known, but it may be a direct consequence of the presence of mutant HTT in myocytes (Orth et al., 2003, van der Burg et al., 2009). Inclusions bodies formed by mutant HTT are present in muscle cells of both HD patients (Ciammola et al., 2006) and mouse models (Moffitt et al., 2009, Sathasivam et al., 1999).

In skeletal muscle, mutant HTT alters the gene expression profile in both mouse models of HD and clinical HD (Luthi-Carter et al., 2002, Strand et al., 2005), involving apoptotic and autophagic pathways and contributing to a catabolic phenotype and muscle wasting (Magnusson-Lind et al., 2014, She et al., 2011).

The R6/2 mouse model is the most commonly used transgenic model of HD that expresses a truncated N-terminal fragment of HTT (Mangiarini et al., 1996), mimicking human HD with central and peripheral pathological changes (She et al., 2011, van der Burg et al., 2008, Magnusson-Lind et al., 2014). The original R6/2 line carried exon 1 of the huntingtin gene with ~150 CAG repeats presented with a rapid and aggressive phenotype similar to juvenile HD (Mangiarini et al., 1996). However, due to the unstable CAG repeat length in breeding practices, the CAG repeat length differs among colonies resulting in a difference in disease severity, age of onset, and life span (Cummings et al., 2012). In general, the HD phenotype is most severe in mice with ~150 CAG repeats, while further increases in CAG length results in milder HD phenotypes (Cummings et al., 2012).

Satellite cells (SCs) are muscle-specific stem cells located between the plasma membrane of the muscle fiber and the basement membrane (Relaix et al., 2021, Mauro, 1961). The stem cell property of satellite cells is essential for their maintenance and is well known in muscle regeneration (Lepper et al., 2011, Sambasivan et al., 2011, Relaix et al., 2021). Satellite cells are normally quiescent but are activated in response to stress induced by injury or exercise. As a result, SCs undergo proliferation prior to differentiation in order to repair and replace damaged myofibers, as well as self-renewal to restore the pool of quiescent satellite cells (Dumont et al., 2015). Previous studies suggest that HD satellite cells undergo a faster rate of programmed cell death and a decreased fusion and regeneration capacity than healthy controls (Ciammola et al., 2006). Satellite cells in HD patients also exhibit less capability of compensating for lost tissue following endurance training (Mueller et al., 2019). Nonetheless, the precise mechanism by which mutant HTT expression in SCs affects muscle tissue formation and regeneration and contributes to overall muscle wasting is not well studied. Several injury models have been exploited to study muscle regeneration kinetics, satellite cell function, and remodeling processes in healthy and disease states, and the most commonly used injury model is intramuscular cardiotoxin (CTX) injection, a toxin derived from snake venom (Hardy et al., 2016). Following an acute injury, the skeletal muscle regeneration process involves a complex process involving satellite cells and activation of innate immune responses including proinflammatory cytokines, such as tumor necrosis factor-α (TNF-α), interleukin (IL)-1 and IL-6 (Yang and Hu, 2018, Baghdadi and Tajbakhsh, 2018). Circulating levels of these proinflammatory cytokines are also elevated peripherally in pre-symptomatic and symptomatic HD patients, as well as mouse models (Bjorkqvist et al., 2008).

The overall hypothesis of this study is that mutant HTT expression in satellite cells leads to abnormalities in both developmental and regeneration processes, which contributes to muscle wasting and motor disturbances in clinical HD. Therefore, we investigated the effects of mutant HTT in satellite cells *in vivo* and *in vitro*. We found that R6/2 neonates have reduced fiber width compared to WT littermates and the reduction in fiber width correlates with R6/2 disease severity *in vitro*. In response to an acute injury, the tissue remodeling process is altered in R6/2 mice, suggesting a satellite cell dysfunction in HD mice.

## Results

### R6/2 neonates exhibit reduced body weight and myofiber width

In this study, we used R6/2 mice from two different colonies expressing different lengths of CAG repeats. In line with previous studies (Cummings et al., 2012), R6/2 mice expressing 266-328 CAG (referred to as R6/2^CAG300^) have a milder HD phenotype than R6/2 mice with 242-257 CAG (referred to as R6/2^CAG250^) repeats. Progressive weight loss is a significant hallmark of mutant HTT gene expression (van der Burg et al., 2008). Our colony of R6/2^CAG300^ mice also exhibited reduced body weight compared to WT littermates at 18 weeks (p= 0.0035) (Figure 1A). We assessed fiber width in longitudinally sectioned and Hematoxylin and eosin (H&E) stained gastrocnemius and soleus muscles in 18-week-old male R6/2^CAG300^ mice and their WT littermates. Stereological quantification showed that in R6/2^CAG300^ mice, both gastrocnemius (p= 0.0307) (Figure 1B) and soleus (p=0.0144) (Figure 1C) muscles had significantly reduced fiber width compared with WT mice. Moreover, we observed increased endomysial connective tissue between R6/2^CAG300^ mice muscle fibers with less connection to adjacent fibers compared to muscle tissue in WT mice (Figure 1D). Next, we stained gastrocnemius muscle sections from 12 weeks old R6/2^CAG250^ male mice and their WT littermates with anti-HTT antibody (em48). In line with previous studies (Orth et al., 2003, Sathasivam et al., 1999), we detected em48 positive inclusions in R6/2^CAG250^ muscle, whereas WT muscle samples were negative for mHTT inclusions (Supplementary Figure 1).

**Figure 1.**
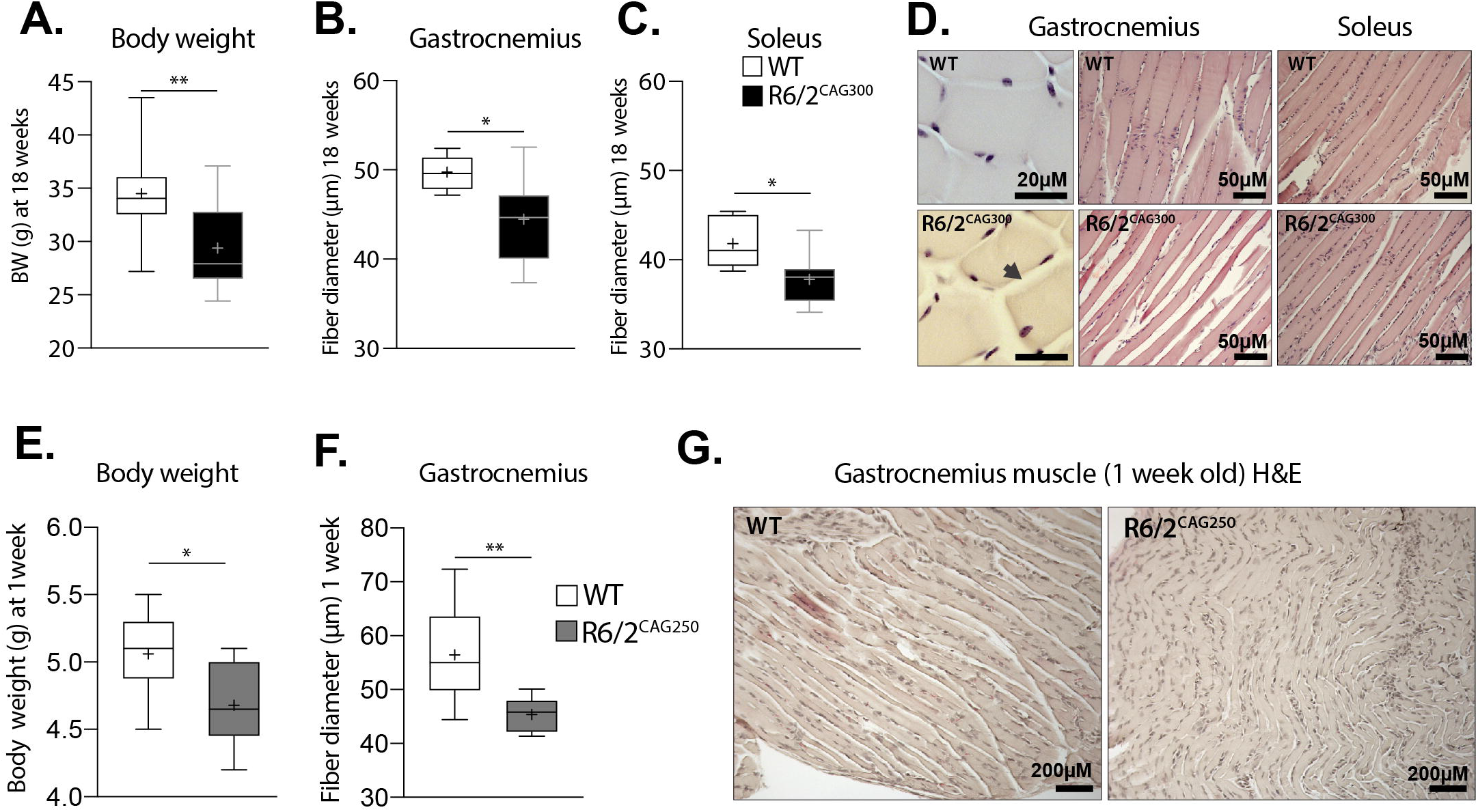
R6/2 mice exhibit reduced body weight and myofiber width. **(A)** Body weight changes in R6/2^CAG300^ and WT mice at 18 weeks (unpaired t-test, n=12-13, p=0.0035). Stereological measurements on H&E stained **(B)** gastrocnemius and **(C)** soleus muscle show reduced fiber diameter in R6/2^CAG300^ mice (unpaired t-test, n=7-8, gastrocnemius: p=0.0307, soleus: p=0.0144) compared to their WT littermates at 18-weeks. **(D)** Representative figures of the gastrocnemius and soleus muscle in 18 weeks old R6/2^CAG300^ mice and their WT littermates. Body weight and gastrocnemius muscle fiber width in 7-9 days old R6/2^CAG250^ WT littermates were assessed. **(E)** The R6/2^CAG250^ have reduced body weight compared to WT littermates at the neonatal stage (unpaired t-test, n=10/group, p=0.0123) and **(F)** reduced gastrocnemius fiber width compared to WT littermates. (Unpaired t-test, n=9-12, p=0.0011). **(G)** Representative figures of the gastrocnemius muscle in 1-week old WT and R6/2^CAG250^ mice. Data are represented as box plots, with boxes representing 25–75 percentile, horizontal lines are median, and whiskers extending to a minimum to maximum values.

Since weight loss and decreased muscle fiber diameter are characteristics of adult R6/2 mice, we further investigated these parameters at the neonatal state. First, we assessed body weight and muscle fiber (gastrocnemius) width in a mixed female and male group of 7-9 days old R6/2^CAG250^ and their WT littermates. Body weight of R6/2^CAG250^ neonates was significantly decreased compared to WT mice (p= 0.0124) already (Figure 1E). Next, we measured gastrocnemius muscle fiber width of R6/2^CAG250^ and WT mice on H&E stained sections. R6/2^CAG250^ neonates exhibited ~16% smaller muscle fiber width compared to their WT littermates (p=0.0011) (Figure 1F and 1G).

### Satellite cells from R6/2^CAG250^ mice exhibit reduced myotube width and altered gene expression *in vitro*

Analysis of changes in whole muscle tissue has the disadvantage of having a smaller number of associated SCs, and the potential effects of mHTT expression in these progenitor cells may occur below the detection limit. Moreover, since the disease severity is milder in mice with ~300 CAG repeats than ~250 CAG repeats, we isolated SCs (7-9 days old neonates) from gastrocnemius from the two R6/2 colonies and assessed their differentiation potentials at 7 days post-differentiation. First, we quantitatively analyzed the myotube width on R6/2^CAG250^, R6/2^CAG300^, and their WT littermates, including both female and male neonates (Figure 2A). We found a significant decrease in myofiber width (~22% decrease) in R6/2^CAG250^ mice compared to WT mice (p=0.0004), whereas there was no difference in R6/2^CAG300^ mice compared to WT mice (p=0.333) (Figure 4A). Yet, the reduction in myotube width in R6/2^CAG250^ group was statistically significant compared to R6/2^CAG300^ mice (p= 0.0097), where R6/2 mice with a shorter CAG repeat length had a ~20% thinner myotube width (One-way ANOVA, Tukey’s multiple comparisons, Genotype F (2, 31) = 9.491, p=0.0006) (Figure 2A and 2B).

**Figure 2.**
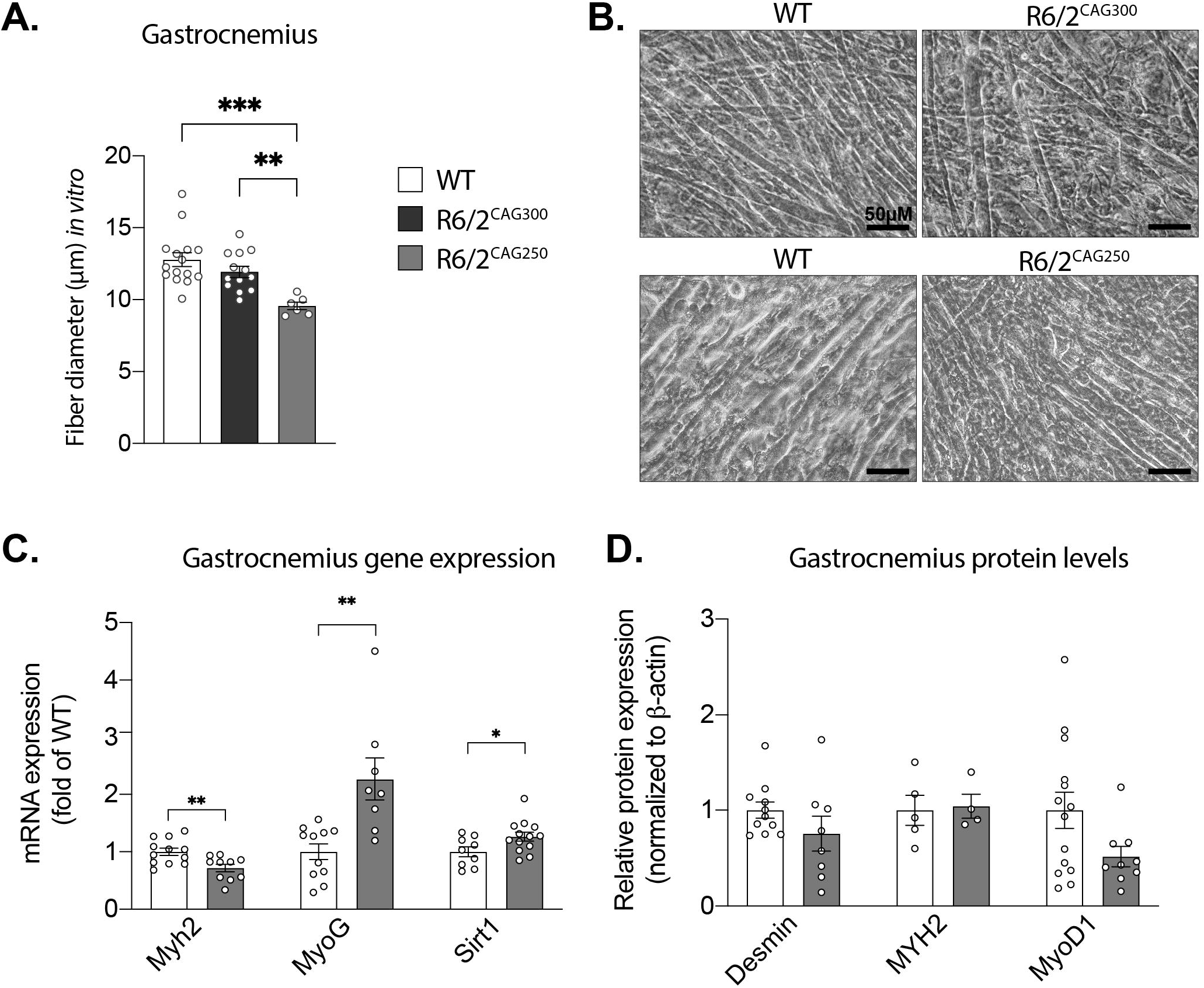
Satellite cells from R6/2^CAG250^ mice exhibit reduced myotube width and altered gene and protein expression *in vitro*. **(A)** Stereological measurements of the fiber width were performed on differentiated satellite cells from R6/2^CAG250^, R6/2^CAG300^, and WT mice at 7 days post-differentiation. The myotube width in R6/2^CAG250^ mice was decreased compared to both R6/2^CAG300^ and WT groups (One-way ANOVA, followed by Tukey’s multiple comparisons test, WT vs. R6/2^CAG250^, p=0.0004 and R6/2^CAG300^ vs. R6/2^CAG250^, p=0.0097). There were no differences in R6/2^CAG300^ mice compared to WT mice (p=0.333). **(B)** Representative images show myotubes from gastrocnemius muscle satellite cells *in vitro*. **(C)** Gene expression analysis in cultured myotubes from R6/2^CAG250^ and WT gastrocnemius muscle. The expression level of *myh2* was reduced in R6/2^CAG250^, whereas *myoG* and *sirt1* were elevated in R6/2^CAG250^ compared to the WT group (the data are represented as fold change in relation to WT cells after normalization to mRNA levels of *actb* and *canx, myh2:* Unpaired t-test, p=0.0060, *myoG:* Unpaired t-test, p=0.0022, *sirt1:* Unpaired t-test, p=0.0366). **(D)** The *in vitro* protein levels of desmin and MYH2 were comparable and there was a trend towards a decrease in MyD1 levels between R6/2^CAG250^ and WT groups at the differentiation day 7. Data represent mean ± SEM.

Previous studies have described alterations in R6/2 mice whole muscle gene expression profile (Strand et al., 2005) (Magnusson-Lind et al., 2014), including activation of apoptotic, HDAC4-Dach2-myogenin axis but also a shift from fast-type fiber types to slow-twitch fibers in tibialis anterior and extensor digitorum longus muscles in HD mouse models (Mielcarek et al., 2015, She et al., 2011). Here, we found a significant increase in *MyoG* (myogenic factor 4) transcription factor involved in myogenesis and repair (p=0.0022) and *Sirt1* (Sirtuin 1) transcription factor and coregulator of several genes (p=0.0366) mRNA levels (Figure 2C) in R6/2^CAG250^. In line with previous studies (Magnusson-Lind et al., 2014), Myosin Heavy Chain 2 (*Myh2*), a fast-twitch and late-state marker (Chal and Pourquie, 2017) gene expression level was significantly decreased (p=0.006) in R6/2^CAG250^ *group in vitro* (Figure 2C).

We next determined early and late-stage myogenesis markers at protein levels, such as desmin, which is essential for the structural integrity and function of muscle (Chal and Pourquie, 2017); MyoD1, which is involved in muscle commitment and differentiation as well as Myh2. We assessed protein levels of these markers in differentiated SCs from R6/2^CAG250^ and their WT littermates *in vitro* at the 7^th^ day of differentiation (Figure 2D). There was no change in early-stage marker desmin (p= 0.1975), which is essential for the structural integrity and function of muscle (Paulin and Li, 2004). The late-stage and fast-twitch marker MYH2 protein levels were unchanged in differentiated SCs from R6/2^CAG250^ (p= 0.9048) compared to the WT group (Figure2D). However, MyoD1 protein level, which is involved in muscle commitment and differentiation, showed a trend toward a decrease in myotubes from R6/2^CAG250^ compared to WT (unpaired t-test, p= 0.0685) (Figure 2D).

### Pax7+ satellite cell pool is reduced in R6/2^CAG300^ mice following acute muscle damage

Normally, SCs are quiescent and become proliferative following an injury, which results in the formation of new and functional muscle fibers (Dumont et al., 2015). To investigate the regeneration capacity of SCs in R6/2 mice, we injected the tibialis anterior muscle of 15 weeks old a mixed group of female and male R6/2^CAG300^ and their WT littermate controls intramuscularly with cardiotoxin or saline (contralateral control muscle). First, we assessed the fiber perimeter on laminin-stained sections at 4 weeks post-injection (Figure 3A). In response to acute injury, the WT-CTX group had a similar fiber area compared to the WT-saline group, and both WT-saline and -CTX groups had significantly wider perimeter than the R6/2^CAG300^-saline group (Figure 3B).

**Figure 3.**
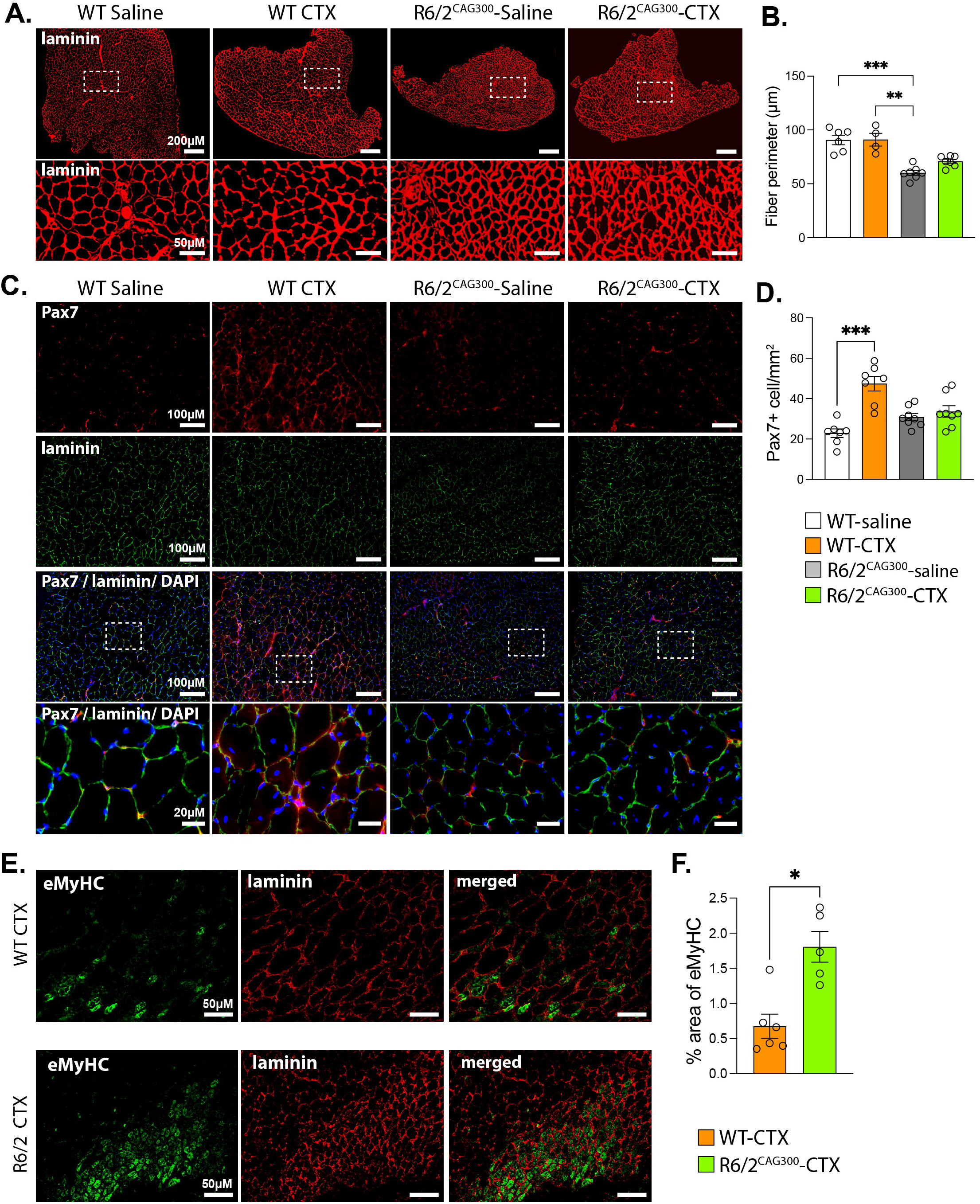
Increased eMHC positive fibers and an unaltered number of Pax7 positive cells in R6/2 compared to WT at 4 weeks post cardiotoxin -injection. **(A)** Laminin immunofluorescent staining showing tibialis anterior muscle from R6/2^CAG300^, and WT mice injected either with saline or cardiotoxin (CTX) at 4 weeks post-injection. **(B)** Tibialis anterior myofiber perimeter of R6/2^CAG300^-saline group is significantly decreased compared to both WT-saline and CTX groups (Kruskal-Wallis test, WT-saline vs. R6/2-saline, p=0.0008 and WT-CTX vs. R6/2^CAG300^-saline, p=0.0029). **(C)** Representative immunofluorescent staining showing the Pax7 positive cells and laminin in saline and CTX injected tibialis muscle at 4 weeks post-injection. **(D)** Quantification of Pax7 positive cells shows an increased number of Pax7+ satellite cells only in CTX injected WT tibialis muscle (Kruskal-Wallis test, WT-saline vs. WT-CTX, p=0.0002). **(E)** Images showing the eMHC positive myofibers and basal lamina in tibialis muscle from R6/2^CAG300^ and WT mice injected either with saline or cardiotoxin (CTX) at 1-week post-injection. **(F)** Percent area assessments of eMHC positive myofiber show that R6/2^CAG300^-CTX mice had a significantly higher level of eMHC expression compared to the WT-CTX group at 1-week post-injection (Mann-Whitney test, p=0173). Data represent mean ± SEM.

To further assess the satellite cell pool following injury, we quantified the Pax7 positive cells at 4 weeks post-injection (Figure 3B). Pax7 is a transcription factor expressed by both active and quiescent SCs; hence it can be used to assess the total number of SCs in a skeletal muscle (Zammit et al., 2006). The number of Pax7 positive SCs was significantly increased in WT-CTX compared to the WT-saline group (p=0.0002), whereas the number of SCs was comparable in R6/2^CAG300^-saline and -CTX tibialis anterior muscles (Figure 3D).

We also assessed embryonic myosin heavy chain (eMyHC), which is skeletal muscle-specific contractile protein and a unique myosin heavy chain isoform expressed during muscle development and regeneration (Agarwal et al., 2020). The R6/2^CAG300^-CTX group had significantly higher levels of eMyHC expressing fibers compared to WT-CTX tibialis anterior muscle at 1-week post-injection (Figure 3E and 3F).

### R6/2^CAG300^ mice exhibit altered inflammatory profile in response to acute damage

Inflammation is a prominent feature of HD muscular wasting, and NF-κB signaling pathway proteins are increased in both HD skeletal muscle and serum (Magnusson-Lind et al., 2014, Sjogren et al., 2017, Bjorkqvist et al., 2008). Therefore, we assessed the inflammatory phenotype using CD68 and CD11b markers in R6/2^CAG300^ muscle in response to acute injury at 1- and 4 weeks post-injection (Figure 4A-4D). The cluster of Differentiation 68 (CD68) protein is highly expressed in macrophages and mononuclear phagocytes. The CD68 positive cell area assessment showed that the percent area of the CD68 cells was increased in both WT-CTX and R6/2^CAG300^-CTX groups compared to only WT-saline control at both 1 week and 4 weeks postinjection (Figure 4E and 4F). We further investigated the percentage area of CD11b positive cells, which are macrophages and microglial cells. The inflammatory CD11b profile of the CTX group was similar to that of the CD68 marker. CD11b positive cells were increased in both WT and R6/2^CAG300^ CTX groups in comparison to WT-saline at both 1 week and 4 weeks postinjection time points (Figure 4G and 4H). In line with other CTX induced acute injury models (Hardy et al., 2016), both CD68 and CD11b responses were increased at 1-week post-injection and dampened later in the regeneration process at 4 weeks post-injection. However, the inflammatory response in R6/2^CAG300^-CTX was not different from the R6/2^CAG300^-saline control group at both time points. Furthermore, a pairwise comparison of WT-saline and R6/2^CAG300^-saline groups showed that at the basal level R6/2^CAG300^ mice had an increase in CD68 and CD11b positive cells (two-tailed Mann-Whitney test, WT-saline vs. R6/2^CAG300^-saline: CD68 at 1-week, p=0.0173, and 4-weeks: p=0043; CD11b at 1-week: p=0.0043, at 4 weeks: p=0.0006).

**Figure 4.**
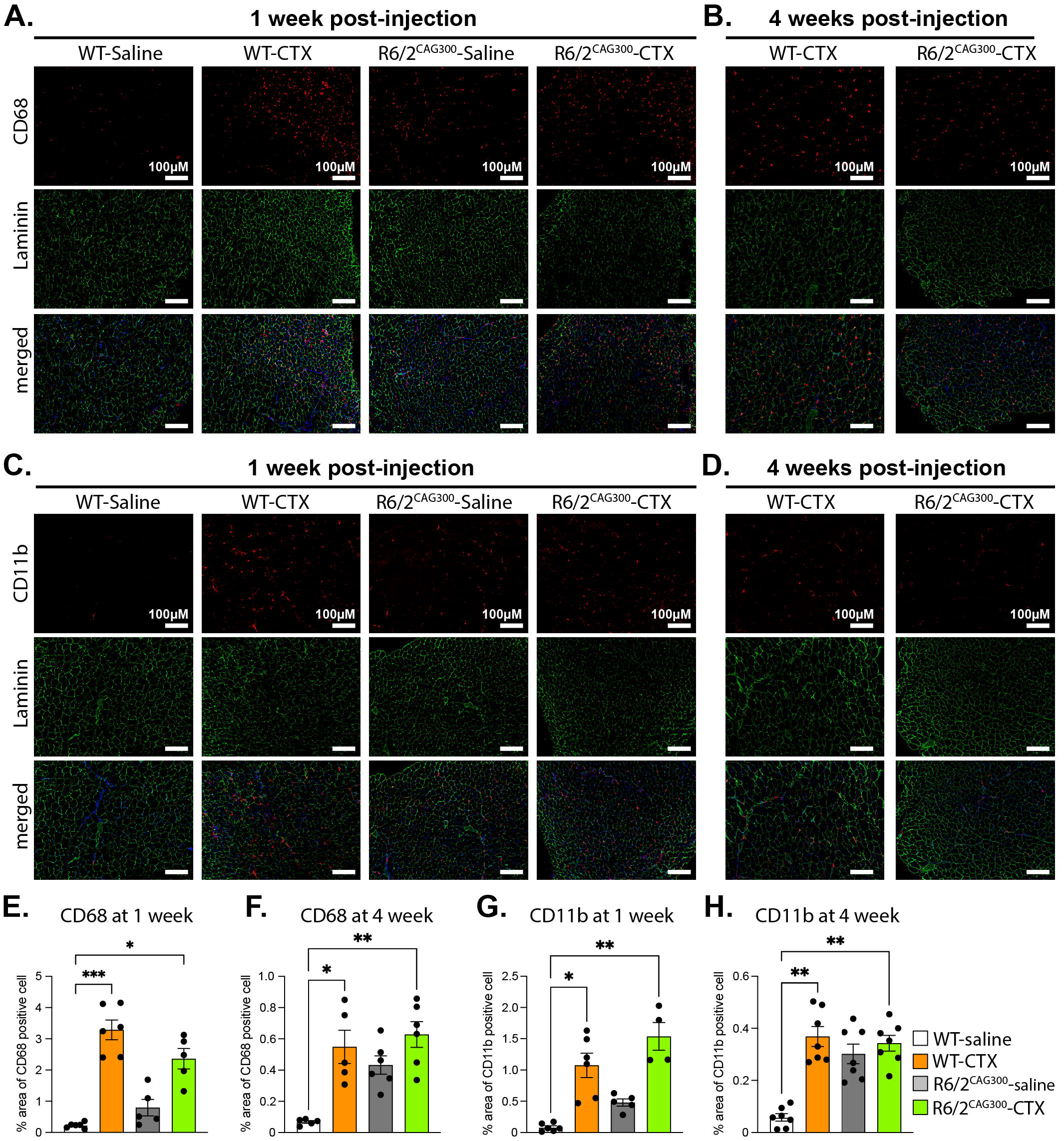
R6/2 skeletal muscle exhibit an altered inflammatory response to acute muscle damage compared to WT littermates. Immunofluorescent staining show CD68 monocyte lineage inflammatory cells and CD11b positive cells at **(A, C)** 1- and **(B, D)** 4-weeks postinjection either with saline or CTX. In response to acute injury, both WT-CTX and R6/2^CAG300^-CTX muscle shows an increase in CD68 cell area compared to only WT-saline group at both **(E)** 1 week (Kruskal-Wallis test, WT-saline vs. WT-CTX, p=0.0005, and WT-saline vs. R6/2^CAG300^-CTX, p=0.0271) and **(F)** 4 weeks (Kruskal-Wallis test, WT-saline vs. WT-CTX, p=0.0444, and WT-saline vs. R6/2^CAG300^-CTX, p=0.0049) post-injection. **(G)** Quantifications of CD11b positive area showed an increase in WT-CTX and R6/2^CAG300^-CTX compared to only WT-saline at both **(G)** 1 week (Kruskal-Wallis test, WT-saline vs. WT-CTX, p=0.011, and WT-saline vs. R6/2^CAG300^-CTX, p=0.0018) and **(H)** 4 weeks (Kruskal-Wallis test, WT-saline vs. WT-CTX, p=0.0016, and WT-saline vs. R6/2^CAG300^-CTX, p=0.0055) post-injection.

## Discussion

In this study, we provide evidence for early pathogenic effects of mHTT on the development of skeletal muscle tissue, *i.e.*, muscle atrophy at the neonatal stage exemplified by reduced skeletal muscle fiber diameter and abnormal tissue morphology. These early changes in R6/2 mice suggest a developmental delay that may underlie reduced muscle strength and atrophy in adulthood.

A neuropathological hallmark for HD are mHTT inclusions observed not only in the brain (Graveland et al., 1985, Vonsattel et al., 1985), but also in muscle and other non-neuronal tissue in both clinical HD (Ciammola et al., 2006) and HD mouse models (Moffitt et al., 2009, Sathasivam et al., 1999). Furthermore, in adult HD skeletal muscle, mutant HTT alters gene expression profiles (Luthi-Carter et al., 2002, Strand et al., 2005), and activates apoptotic and autophagic pathways and eventually contributing to severe muscle wasting at the end-stage (Magnusson-Lind et al., 2014, She et al., 2011). In line with previous studies, the signs of muscle atrophy were evident in adult R6/2 mice, and it is mainly attributed to degenerative effects of mHTT over the disease progression. However, we found that fiber width is also significantly reduced in 1-week old R6/2^CAG250^ neonates and cultured R6/2^CAG250^ myotubes, indicating that pathogenic developmental effects of mHTT could be one of the main contributing factors of HD skeletal muscle pathogenesis.

Even though toxic or preventive effects of inclusions are controversial (Arrasate et al., 2004, Kim et al., 1999, Sakahira et al., 2002), in line with previous studies (Sathasivam et al., 1999), we detected intranuclear mHTT inclusions 12 weeks old R6/2^CAG250^ gastrocnemius muscle, and we observed that a fraction of nuclei were positive for mHTT inclusions. The subcellular localization of mHTT inclusions should be further defined to understand whether myonuclei or quiescent SCs are positive for inclusion bodies. However, this inhomogeneous distribution of inclusions could also be due to high muscle turnover in R6/2 (newly formed myotubes or degenerating mature myotube) or differences in ubiquitin-proteasome system and autophagic activity of muscle fibers (Orth et al., 2003).

Satellite cells, progenitors of mature skeletal muscle fibers (Relaix et al., 2021) (Mauro, 1961), are essential for the proliferation, differentiation, and repair of muscle tissue in response to muscle injury or exercise (Dumont et al., 2015). Satellite cell function and renewal capacity are affected in several muscular dystrophies such as Duchenne muscular dystrophy (Ervasti et al., 1990), where SCs lose the ability to establish asymmetric satellite stem-cell divisions and cell polarity that leads to severe muscle wasting by causing myofiber instability and impaired regeneration (Chang et al., 2016). Since mutant HTT is ubiquitously expressed in the body, it can also have a direct pathological effect on the SCs (Chang et al., 2016) and HD muscle wasting could be the direct consequence of mutant HTT presence in SCs. To test our hypothesis, we assessed the SCs’ functional capacity in R6/2 mice with two different CAG repeat lengths (R6/2^CAG250^ and R6/2^CAG300^) *in vitro*. Our colonies’ long CAG repeats size results in a slower disease progression than the original R6/2 mice with 150 CAG repeats (Morton et al., 2009). A slower disease progression in R6/2 mice with longer CAG repeats provide a broader window to evaluate HD pathologies and resemble more adult-onset HD phenotype regarding the phenotype severity and disease onset. As a result, R6/2^CAG250^ mice exhibit a more severe HD phenotype compared to R6/2^CAG300^ mice (Cummings et al., 2012). And in line with this, we found that differentiated SCs from R6/2^CAG250^ colony formed thinner myotubes compared to R6/2^CAG300^, which validates the R6/2 phenotypical HD pathology.

Mitochondrial dysfunction has been suggested as a critical player in skeletal muscle pathology in HD, with impaired PGC-1α function (Chaturvedi et al., 2009). Sirt1 has been suggested to act as a sensor of energy deprivation and regulate metabolism homeostasis by activating PGC-1α (Canto and Auwerx, 2009). Increased expression of *Sirt1* has been shown to impair muscle differentiation both *in vitro* (Fulco et al., 2003) and *in vivo* (Toledo et al., 2011, Sadagurski et al., 2011), resulting in muscle repair deficits. In this study, we found increased levels of *Sirt1* mRNA levels in cultured SCs from R6/2^CAG250^ gastrocnemius muscle compared to WT littermates. This indicates that energy deprivation alongside decreased regeneration ability is also present *in vitro*.

MyoD knock-out skeletal muscle exhibits an elevated number of satellite cells with a lesser extent of muscle differentiation competency, and in the case of injury, lack of MyoD reduces the regenerative capacity. At the protein level, we found reduced myotube width and a trend towards diminished MyoD expression in differentiated SCs from R6/2^CAG250^ compared with WT littermates *in vitro*. Considering these skeletal muscle lineage determination functions of MyoD, decreased protein levels of MyoD could be responsible for the blunted myotube formation capacity found in the R6/2^CAG250^ model of HD.

SCs proliferate under any critical conditions (*i.e.,* mechanical or chemical damage) to form new fibers to recover from muscle atrophy in healthy conditions (Relaix et al., 2021). Therefore, using a chemical injury-induced approach, we investigated muscle regeneration capacity and satellite cell pool of R6/2 mice *in vivo*. Following any acute muscle damage, it is known that WT muscle reorganization appears histologically normal with an increased number of Pax7 positive SCs at 4 weeks post-injection (Hardy et al., 2016). Yet, the R6/2^CAG300^-CTX and R6/2^CAG300^-saline groups had a comparable level of Pax7 positive cells, lacking a normal response to tissue remodeling. Furthermore, as R6/2^CAG300^-CTX muscle also had a significantly higher level of eMyHC positive cells compared to the WT-CTX group at 1-week post-injection, a group of Pax7+ cells might still be committed to differentiation, indicating delayed regeneration process or depleted satellite cell pool in HD mice. However, this should be further investigated by assessing both Pax7/MyoD1 positive cells at 4 weeks post-injection and levels of eMyHC at a later point than 1-week post-injection.

In response to CTX, an immune response is triggered in a normal muscle (Tidball, 2017). Indeed, we found an increase in CD11 and CD68 positive cells in WT muscle 1 week after CTX injection. In HD, activation of the NF-κB signaling pathway with elevated levels of p65/RelA, Tradd, and TRAF5 has been shown in both R6/2 and full-length knock-in HdhQ175 mouse models (Magnusson-Lind et al., 2014, Sjogren et al., 2017). In this study, we found a trend towards an increase in CD68 and CD11b positive cells in the R6/2^CAG300^-saline group compared to WT mice. Possibly, this elevated basal inflammatory state in R6/2^CAG300^ skeletal muscle is related to the lack of immune response to damage caused by CTX injection in the R6/2^CAG300^-CTX group.

To understand the developmental effects of mutant HTT expression in skeletal muscle, the satellite cell renewal, and myogenic commitment processes should be further investigated at the gene expression and protein levels. As membrane permeability to Ca^2+^ ions could be affected in HD (Kolobkova et al., 2017) and the elevated concentration of Ca^2+^ could lead to muscle dystrophy (Millay et al., 2009, Turner et al., 1991), the Ca^2+^ status of myotubes should be further investigated using Ca^2+^ imaging techniques.

---

**Supplementary figure 1: em48 positive inclusions in muscle fiber nuclei of R6/2 mouse models.** Representative pictures of skeletal muscle showing mutant HTT inclusions 12 weeks old R6/2^CAG250^ male mice (n=3) gastrocnemius muscle and their WT littermates (control). Gastrocnemius muscle sections were double-stained for em48 and Laminin to visualize inclusions and basal lamina of myofibers, and the nucleus was visualized using DAPI. We detected em48 positive inclusions in R6/2^CAG250^ group, while WT gastrocnemius muscle sections were negative for the inclusions.

## Materials and Methods

### Ethics

Animal experiments were carried out in strict accordance with Swedish legislation and approved by local and national regulatory authorities (Animal Ethics Committee in Lund and Malmö, Sweden; ethical permit numbers: 10992/18 and 07456/21).

### Animals

Transgenic R6/2 HD mice (expressing exon 1 of the HD gene) (Mangiarini et al., 1996) on the C57BL/6xCBA genetic background (Jackson Laboratory, Bar Harbor, ME, USA) and their wild-type (WT) littermates were used in this study. The CAG-repeat lengths of the R6/2 mice from two different colonies used in this study ranged either between 242-257, and 266-328. Genotyping of the R6/2 colony was performed on ear punch samples for the R6/2 gene using PCR. The PCR product was separated by gel electrophoresis using a 2% agarose gel for detecting the R6/2 gene. All primers were purchased from MWG Eurofins (Eurofins MWG Synthesis GmbH, Germany), and sequences of primers used for R6/2 colony genotyping in this study are found in Supplementary Table 1. The CAG repeat size of our colony was obtained from tail biopsies using polymerase chain reaction assay by Laragen (Laragen Inc., CA, USA). Two separate R6/2 mice colonies had CAG repeat length of 242-257 (R6/2^CAG250^), and 266-328 (R6/2^CAG300^) repeats, all have a slower disease progression than the original R6/2 mouse line with 150 CAG repeats, as described by Morton and co-workers in 2009 (Morton et al., 2009). The mice were housed in groups of 2-5 in universal Innocage mouse cages (InnoVive, San Diego, CA, USA) under standard conditions (12 h light/dark cycle, 22°C) ad libitum access to standard chow and water.

### Cardiotoxin injections

The 15 weeks old R6/2^CAG300^ and WT mice were anesthetized using isoflurane and placed in a prone position. Cardiotoxin (*Naja pallida*, Sigma #217503) was prepared in saline solution and 30 μl of 10 μM cardiotoxin was injected into the anterior compartment and the same volume of saline solution was injected into the contralateral compartment. The R6/2^CAG300^ and WT mice were deeply anesthetized using pentobarbital and the tibialis anterior muscle were harvested on post-injection days 7, and 28. The muscles were embedded in OCT and slowly frozen using liquid nitrogen. The fresh frozen tissue was sectioned at 7 μm, the specimen was placed on plus coated slides.

### Cell culture preparation

Gastrocnemius muscle from 7-9 days old R6/2^CAG250^, R6/2^CAG300^ and WT mice were used for the *in vitro* assessment of satellite cells in HD. The mice were euthanized by decapitation, and the gastrocnemius muscle from both left and right back limbs was dissected. An incision was made using a scalpel from the heel to the tail line. First, the skin and then the muscle epimysium was removed to expose the gastrocnemius muscle. The Achilles tendon of gastrocnemius was cut, and the adjacent soleus muscle was removed. The whole gastrocnemius muscle was dissected by releasing it from the femur bone and transferred to a falcon tube with 1ml DMEM and kept on ice. Using blades, the gastrocnemius muscle was cut into small pieces and minced into a smooth pulp. To dissociate the muscle, the muscle was then transferred to a 15 ml Falcon tube with 0.5% collagenase/dispase and placed in a water bath at 37°C and shaken gently every 10 minutes until cell suspension was obtained. The cell suspension was triturated gently to homogenize the mixture and to dissociate it into a single-cell suspension. 2% FBS in PBS up to 9 ml was added to the single-cell suspension and mixed well. A cell strainer (40 μm pore size) was placed onto a 50 ml Falcon tube and the cell suspension was transferred onto the cell strainer. The strainer was rinsed with an additional 5 ml 2% FBS in PBS. By centrifuging the tubes at 500G at 21 °C for 5 min, mononuclear cells were collected. Following that, mononuclear cells were plated on a Matrigel-coated 12-well plate. The cells were left for proliferation for 3 days with 1 ml of proliferation media containing Ham’s 12, 20% FBS, 20mM L-glutamine, 1%penicilin/streptomycin +1% amphotericin B. The plates were kept in a 5% CO_2_ incubator at 37°C. To induce differentiation of satellite cells, growth media was replaced with differentiation media containing Dulbecco’s Modified Eagle Medium (DMEM), 2% ml horse serum, 1% penicillin/streptomycin and 1% amphotericin B. The differentiation media was refreshed every day and incubated for 7 days. For further assessment of the myotube width, 25 pictures at magnification 20x were taken randomly at 5 locations for each well. For protein extraction of differentiated satellite cells, the differentiation media was replaced with 250 μL cold 2% SDS lysis buffer and collected using a Fisherbrand cell scraper and carefully transferred to 1.5 mL Eppendorf tubes for further protein extraction assessments.

### Assessment of myotube width *in vitro*

The 25 pictures taken at magnification 20X of the myotubes after differentiation was used for stereological analysis, using ImageJ 1.50 software (National Institutes of Health, Bethesda, MD). Using a straight-line probe, the myotube width was assessed, and the results were obtained in the ROI manager. The myotubes were measured in three locations, top, middle, and bottom, length was calculated in the perimeter and 600-900 measurements were done per genotype.

### Histology of skeletal muscle *in vivo*

Skeletal muscle gastrocnemius and soleus were dissected, placed into 4% paraformaldehyde in phosphate-buffered saline (0.01 M) overnight and then moved into 70% ethanol until paraffin embedding for further stereological assessment. Sections (7 μm) were mounted on glass slides and stained according to Mayer’s Hematoxylin-Eosin staining protocol. Digital images of the stained sections were used to identify the morphological features of the muscle. A bright-light microscope (Olympus U-HSCBM, Olympus, Tokyo, Japan) with a 20x or 40x magnification objective, digital camera, and image capture software (cellSens Dimensions 1.11 software; Olympus, Tokyo, Japan) were used. The stereological measurements were performed under blinded conditions.

### Immunofluorescence analysis

In order to visualize mHTT inclusions in skeletal muscle fibers, paraffin-embedded sections (7μm) of skeletal muscle gastrocnemius from 12 weeks old WT and R6/2^CAG250^ male mice were stained for HTT (em48), Laminin and DAPI. Deparaffinized sections went through antigen retrieval using 0.01M citrate buffer for 20 min at 95°C before pre-incubation with 5% normal donkey serum (NDS), 0.25% Tx, 1% BSA, and Tris HCl pH 6 for 1 hour at RT. The sections were incubated overnight at 4°C in a pre-incubation solution containing mouse monoclonal em48 (Merck Millipore; MAB5374, 1:400) and rabbit polyclonal Laminin (Abcam, ab11575, 1:500) antibodies. The CD68 primary antibody incubation was performed at room temperature for 2 hours (mouse CD68, BIO-RAD, MCA1957, 1:750) and CD11b staining (BD Pharmigen, 550282, 1:250), sections were incubated overnight at 4°C. Following the primary antibody incubation, sections were washed three times with PBS. Next, the sections were incubated for 2 hours in pre-incubation solution containing secondary Donkey AF488 anti-mouse (Jackson

ImmunoResearch, 715-545-150, 1:500) or Donkey Cy3 anti-rabbit (Jackson ImmunoResearch; 711-165-152, 1:500) and DAPI. The sections were imaged using a fluorescence microscope (Olympus U-HSCBM, Olympus, Tokyo, Japan) with a 20x magnification objective (cellSens Dimensions 1.11 software; Olympus, Tokyo, Japan).

For MHC and Pax7 staining, 7-μm thick sections were fixed with 4% paraformaldehyde in PBS at room temperature for 5 min and permeabilized with pre-cooled methanol at −20°C for 5 min. After washing with PBS, sections went through antigen retrieval using 0.01M citrate buffer for 5 min at 80°C and another 5 min at 90°C. Any nonspecific antibody binding sites were blocked by incubation in 10% normal donkey serum (for MHC) and 5% bovine serum albumin (for pax7) in PBS at room temperature for 3 hours. Next, they were incubated with mouse fab (0.05mg/mL) (Jackson ImmunoResearch, 015-000-007) for 1 hour at room temperature. Then sections were incubated in primary antibody solutions, *i.e*., embryonic mouse myosin heavy chain (F1.652, DSHB; 1:200) overnight at 4°C and mouse Pax7 (DSHB, 1:1000) for 2 hr at RT. Afterward, cryosections were incubated with a pre-incubation solution containing appropriate secondary antibodies. For MHC, sections incubated with Donkey AF488 anti-mouse (Jackson ImmunoResearch, 715-545-150, 1:1000) for 45 min and for Pax7, Biotin-conjugated Goat Antimouse (Jackson ImmunoResearch, 115-065-205, 1:1000) and Donkey AF488 anti-rabbit (Invitrogen, A-21206, 1:1000) were used to incubate the sections for 45 min and stained with AF568 streptavidin (Invitrogen, S11226) for 30 min at RT.

### Image analysis

Sections were imaged using a fluorescence microscope (Olympus U-HSCBM, Olympus, Tokyo, Japan) with a 4x or 10x or 20x magnification objective (cellSens Dimensions 1.11 software; Olympus, Tokyo, Japan) and all images were analyzed using ImageJ software.

In laminin staining, 4-7 animals were used per group, and at least two sections were imaged at 4x magnification. To assess myofiber perimeter, Trainable Weka Segmentation plugin was used. To assess the percent area of CD68, CD11b, and eMHC and the number of Pax7 positive cells, three images at 10x magnification were taken from 1-2 muscle sections per animal. Using ImageJ built-in plugin, three images from each section were stitched, and percent areas were assessed. For Pax7 quantification, at least one section was imaged from each animal. Then Pax7 positive cells were manually counted using ImageJ cell counter probe. All the analyses were done under blinded conditions.

### RNA extraction and cDNA synthesis

The E.Z.N.A. Total RNA Kit II (Omega bio-tek, Norcross, Georgia, USA) was used to extract total RNA from cell culture on the 7^th^ day of differentiation. The complementary DNA (cDNA) was synthesized using the iScript cDNA Synthesis Kit (Bio-Rad Laboratories, CA, USA), according to the manufacturer’s protocol. RNA concentration and purity were measured by a NanoDrop Lite spectrophotometer (Thermo Fisher Scientific, Wilmington, Delaware, USA).

### Real-time quantitative PCR

SsoAdvanced Universal SYBR Green Supermix from Bio-Rad Laboratories was used for RT-qPCR and performed following the manufacturer’s instructions. All RT-qPCR plates were run on a CFX96 Touch real-time PCR detection system (Bio-Rad, CA, USA). Primers utilized for RT-qPCR validations (see Supplementary Table 2) were designed using either QuantPrime69 or PrimerQuest from Integrated DNA Technologies (http://eu.idtdna.com/PrimerQuest). The efficiency of each primer pair was tested before use by performing a standard curve, and the efficiency criteria for using a primer pair was 90% < E < 110%, with an R^2^ cut-off >0.990. Housekeeping genes, β-actin (*actb*) and ATP synthase, H+ transporting mitochondrial F1 complex, beta subunit (*Atp5b*) were used for normalization. Changes in gene expression were calculated using the CFX manager software program (Bio-Rad, CA, USA), using the ΔΔCt method with a fold change cut-off at ≥1.5 and p < 0.05 considered significant. All samples were run in triplicate, and relevant positive and negative controls were run on each plate.

### Protein extraction

Proteins from differentiated satellite cells (DD7) were extracted using 250 μl lysis buffer containing 2% SDS, 0.1 M Tris-HCl pH 7.4, 50 mM NaCl, 1 mM EDTA, supplemented with protease inhibitors (Complete, Roche, Basel, Switzerland) and phosphatase inhibitors (PhosSTOP, Roche, Basel, Switzerland). The homogenate was incubated in the cold room rolling for 30 min and centrifuged every 10 min at 12000 rpm for 1 min at 8°C. The last centrifugation was for 20 minutes at 12000 rpm at 4°C. The supernatant was collected, and the Pierce BCA Protein Assay kit was used to measure the protein concentration.

### Western blot

20 μg of protein was mixed with 4× Laemmli Sample buffer complemented with 10% β-mercaptoethanol, before denaturing at 95 °C for 10 min. Lysates were run on 4-15% Mini-PROTEAN TGX Precast Gels (Bio-Rad, CA, USA) and blotted on 0.2 μm Trans-Blot Turbo Midi Nitrocellulose membrane (BioRad CA, USA), which were blocked with 5% non-fat dry milk in Tris-buffered saline, pH 7.6, containing 0.1% Tween20 (TBS-T) for 2 hrs at room temperature (RT). Blocked membranes were incubated with rabbit polyclonal Desmin (Abcam, ab15200, 1:500), mouse monoclonal MyoD1 [5.2F] (Abcam, ab16148, 1:500), mouse monoclonal Fast Myosin Skeletal Heavy chain [MY-32] (Abcam, ab7784, 1:500), antibody in 5% non-fat dry milk in TBS-T overnight at 4 °C. After washing several times with TBS-T, membranes were incubated for 2 hrs with a secondary Donkey Anti-Rabbit IgG H&L (Abcam; ab16284, 1:10000) or Donkey Anti-Mouse IgG H&L (Abcam, ab6820, 1:10000) horseradish peroxidase-conjugated antibody. Signal was visualized using Western Blotting Luminol Reagent (Santa Cruz, TX, USA) and imaged using a ChemiDoc MP Imaging System (Bio-Rad, CA, USA). B-actin was used as a loading control and for protein normalization using ImageJ 1.50 software (National Institutes of Health, Bethesda, MD).

### Statistical analyses

The data analysis was performed using GraphPad Prism (GraphPad Software Inc., version 8.01, San Diego, CA, USA). Shapiro-Wilk normality test was used to determine a Gaussian distribution. One-way or two-way factor analysis of variance (ANOVA) with Tukey’s post hoc test was used for multiple comparisons, Kruskal-Wallis with Dunn’s post-hoc test or two-tailed Student’s t-test was used for comparisons. Results are presented as means ± SEM. Differences with a p<0.05 were considered statistically significant.

## Contribution statement

R. S.K, M.B., M.D.H., V.A., K.G., M.S. and S.H. conceived and designed the experiments. S. H., M.S., V.A., N.F., K-G. and R.S.K. performed the experiments and analyzed the data. R.S.K., M.B., S.H., and M.S. wrote the first draft of the manuscript. All authors reviewed the manuscript and approved the final version.

## Acknowledgments

We thank Dr. Bruno Cadot (Sorbonne University-Inserm, Research Center in Myology, Paris, France) for the Weka segmentation plugin and Catarina Blennow, Alicja Flasch, Ann-Charlotte Selberg and Susanne Jonsson for valuable technical assistance.

