## Supplementary figures and images for "Skeletal muscle regeneration is altered in the R6/2 mouse model of Huntington’s disease"

### Supplemental Figure 1

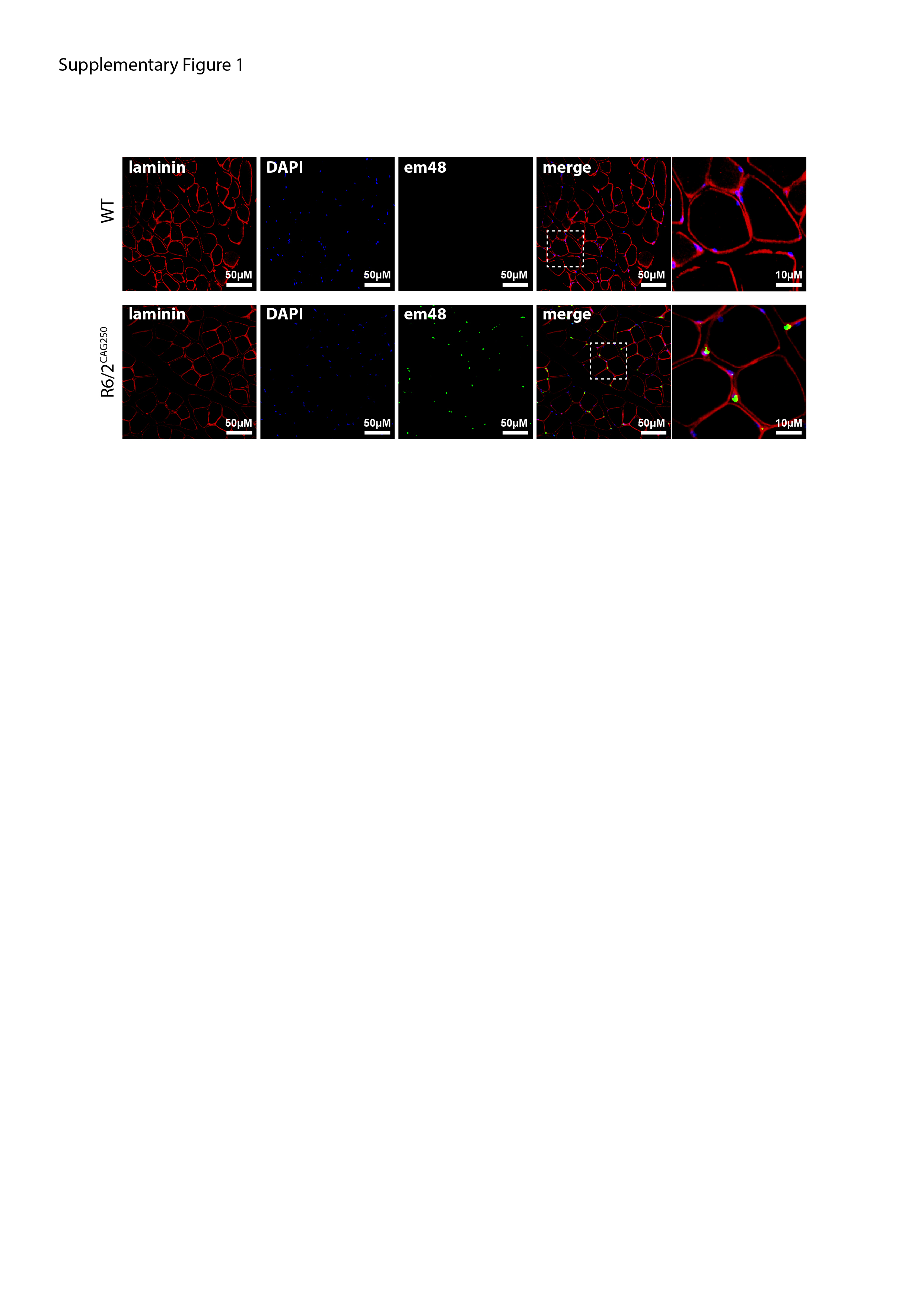
